# Structure-guided computational design and mechanistic understanding of the p95HER2-targeting NAZ-mAb antibody and its variants

**DOI:** 10.64898/2026.07.07.736817

**Authors:** Puneet Rawat, Jon Amund Kyte, Victor Greiff, Esmaeil Dorraji

## Abstract

Human epidermal growth factor receptor 2 (HER2) is an oncogenic receptor tyrosine kinase in breast cancer and other malignancies. A subset of HER2-positive tumours expresses 611-CTF-p95HER2, a tumour-specific, hyperactive truncated isoform associated with metastasis and treatment resistance that lacks most of the extracellular domain targeted by conventional HER2-directed antibodies. We previously developed NAZ-mAb (formerly known as Oslo-2), a monoclonal antibody against 611-CTF-p95HER2. Here, we describe a computational antibody-engineering workflow for designing variants of NAZ-mAb. Starting from the sequence alone, we modeled the NAZ-mAb–611-CTF-p95HER2 complex, generated a combinatorial mutational landscape using FoldX 5.0, and prioritized candidate variants using predicted interaction energy and developability criteria. Two variants representing distinct design strategies were selected for validation: an aromatic double mutant, NAZ-mAb v1 (L:S31W/L:H107W), and a conservative single mutant, NAZ-mAb v2 (L:S31M). Both variants were successfully expressed as recombinant IgGs; NAZ-mAb v2 achieved a five-fold higher recombinant expression yield than parental NAZ-mAb, while both variants retained antigen binding with a higher apparent signal than the parental antibody in indirect ELISA. However, Biacore two-state kinetic analysis revealed weaker affinities than the parental antibody (K_D_ NAZ-mAb v1: 32.6 nM, NAZ-mAb v2: 9.45 nM vs. parental NAZ-mAb: 5.33 nM). These findings show that the computational workflow can generate experimentally tractable, antigen-engaging NAZ-mAb variants, while also highlighting the limitations of fixed-backbone interaction-energy ranking as a predictor of binding affinity and yield. This study provides a practical framework for computationally driven, developability-aware antibody optimization in the absence of experimental structural data.

## 1. Introduction

Breast cancer is the most frequently diagnosed cancer in women, with an estimated 2.3 million new cases and 670,000 deaths globally in 2022 (Kim et al. 2025), and it accounts for nearly one-third of new cancers in women in the United States (Siegel et al. 2026). It is a highly heterogeneous disease encompassing biologically and clinically distinct subtypes with differing molecular drivers, prognosis, and treatment responses (Caswell-Jin et al. 2021). Human epidermal growth factor receptor 2 (HER2) is amplified in roughly one-fifth of breast cancers and is the target of trastuzumab and related biologics (Owens et al. 2004). HER2 carboxy-terminal fragments (CTFs), collectively termed p95HER2, can arise through more than one mechanism. Critically, the hyperactive 611-CTF isoform is generated by alternative translation initiation at the internal AUG encoding methionine 611, rather than by proteolytic shedding of full-length HER2 (Anido et al. 2006). 611-CTF retains the transmembrane and intracellular kinase domains but lacks most of the HER2 extracellular domain, creating a constitutively active receptor that promotes mammary tumour growth and metastasis in experimental models (Pedersen et al. 2009). Clinically, a high p95HER2/HER2 ratio has been associated with inferior progression-free and overall survival in trastuzumab-treated HER2-positive metastatic breast cancer (Chumsri et al. 2018; Sperinde et al. 2010). Because 611-CTF lacks the extracellular regions recognised by trastuzumab and pertuzumab, the truncated receptor is not directly engaged by these conventional HER2-directed antibodies (Scaltriti et al. 2007). Importantly, alternative translation initiation at M611 creates a distinct N-terminal extracellular context that can be selectively targeted. Although the underlying amino-acid sequence is present within full-length HER2, the N-terminal epitope exposed in 611-CTF is not accessible in the intact receptor. We previously developed NAZ-mAb, which selectively recognises 611-CTF-p95HER2 over full-length HER2 and maps to the N-terminal epitope PIWKFPDEE (Dorraji et al. 2022). NAZ-mAb therefore provides a molecularly selective targeting scaffold for the 611-CTF-p95HER2 isoform and is the engineering substrate for the present study.

Mechanistic understanding of binding behaviour, and rational improvement of an antibody, requires a structural model of the paratope–epitope interface, yet experimental structures of antibody–antigen complexes remain scarce due to technical difficulty and high cost. Recent advances across computational structural biology now make it feasible to address this gap through hybrid computational-experimental approaches at reduced cost and time. Sequence-based structure prediction yields high-quality models of both antibody and antigens (Jumper et al. 2021; Leem et al. 2016), but prediction of the antibody–antigen complex itself lags behind (Smorodina et al. 2026). A recent benchmark study reported that more than half of the antibody–antigen complex predictions failed to reach acceptable accuracy (Xu et al. 2025), largely because the hypervariable CDR-H3 loop and the absence of co-evolutionary signal at the interface are poorly captured.

Prior knowledge of the epitope can substantially improve docking, and such information can be obtained from comparatively inexpensive experimental approaches (site-directed mutagenesis, overlapping-peptide scanning or hydrogen–deuterium-exchange mass spectrometry) rather than from structure-determination techniques (such as X-ray crystallography or cryo-EM). This restraint-guided, consensus modelling can potentially improve the antibody–antigen complex structure prediction for downstream analysis. A therapeutic antibody must satisfy two objectives simultaneously: high, specific affinity and good developability (stability, low aggregation and manufacturability). These objectives frequently conflict and must be balanced when designing new variants (Raybould et al. 2024; Makowski et al. 2022). Although many optimisation methods exist, current machine-learning approaches are constrained by limited training-set size and poor interpretability (Akbar et al. 2022). We therefore deliberately used interpretable, physics-based methods for affinity and developability assessment, so that the basis of each design decision can be understood mechanistically, and validated against recombinant expression, indirect ELISA and surface-plasmon-resonance measurements. The central value of the study is mechanistic: aggressive mutagenesis of the parental antibody produced larger and less predictable changes, whereas conservative substitutions yielded only incremental effects, and the most robust strategy proved to be conservative substitution at positions not directly involved in antigen contact. We further show that empirical, fixed-backbone energy functions are systematically biased toward aromatic substitutions, which are over-rewarded through favourable packing and π-contacts despite their conformational and desolvation costs. These insights are transferable to other antibody–antigen systems where experimental structural information is lacking.

## 2. Materials and Methods

### 2.1 Complex modelling and interface definition

The extracellular domain of 611-CTF-p95HER2 was modeled using AlphaFold 2.0 (Jumper et al. 2021) and the NAZ-mAb antibody scFv (and variant) structure with AbodyBuilder (Leem et al. 2016; Abanades et al. 2023). Three docking algorithms, Zdock 3.0.2 (Pierce et al. 2014), Haddock 2.4 (Honorato et al. 2024), and ClusPro 2.0 (Brenke et al. 2012) were used in information-driven docking to predict the top 10 docking poses. The experimentally-validated “MPIWKFPDEE” region in the extracellular domain of p95HER2 was provided as a potential epitope region for docking (Dorraji et al. 2022). The final complex was selected by consensus across the three methods, scored by the percentage epitope overlap.

### 2.2 Mutational scanning and combinatorial modelling

FoldX 5.0 (Delgado et al. 2019) was used to generate the mutant structures and to compute interaction energy and stability. The wild-type complex was first relaxed with the “RepairPDB” command. Paratope and epitope residues were defined by an inter-chain heavy-atom contact distance cut-off of 4.5 Å. Complete single-point saturation mutagenesis (19 substitutions per position) was performed across all paratope residues with “BuildModel” command. The interaction energy was calculated with the “AnalyseComplex” command for wild-type and each mutant structure. The change in interaction energy (Δ Interaction energy) with respect to wild-type was calculated as:

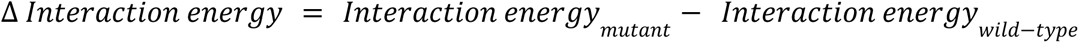

Where, a negative change in interaction energy indicates improved binding affinity. Positions at which more than half of the 19 substitutions improved interaction energy were defined as binding hotspots. To limit the combinatorial multi-site mutation space, only high-confidence binding-improving substitutions (ΔΔG_interaction < −1.5 kcal/mol, well beyond the typical error of empirical ΔΔG prediction) were retained. An exhaustive combination of up to six substitutions produced 35,279 unique multi-site mutants, each scored for change in interaction energy relative to wild-type. All positions or mutations are defined as [chain]:[wild-type][position][mutation]. The stability of the complex was calculated using the “Stability” command in FoldX 5.0 and using DynaMut2 server (Rodrigues et al. 2021). The interaction energy was also validated using MutaBind2 server (Zhang et al. 2020). The structures of the variants generated by FoldX 5.0 were aligned to the parental NAZ-mAb in PyMOL 3.1 (https://www.pymol.org/).

### 2.3 Developability and aggregation profiling

Developability of selected mutants was assessed with the Therapeutic Antibody Profiler (TAP) (Raybould et al. 2024), reporting total CDR length, CDR-vicinity patches of surface hydrophobicity (PSH), positive and negative charge patches (PPC/PNC), and the structural Fv charge symmetry parameter (SFvCSP), each interpreted against the published amber/red guideline thresholds (as of June 2026). Aggregation-prone regions (APRs) were predicted with ANuPP (Prabakaran et al. 2021) using the default model threshold.

### 2.4 Gene synthesis, expression and purification

Variable heavy and light domains for the selected NAZ-mAb antibody variant were codon-optimised for human-cell expression, synthesised with cloning restriction sites, and inserted into mouse-IgG1 heavy- and light-chain expression vectors. Sequence identity was further verified by Sanger sequencing. HEK293 cells were passaged to an appropriate stage for transient transfection. Heavy-chain and light-chain expression vectors were co-transfected, and cells were cultured for approximately 6 days before harvest. Cultures were clarified by centrifugation and filtration. Antibodies were purified by Protein A affinity chromatography, eluted using citrate buffer at pH 3.0, neutralized with Tris buffer, and buffer-exchanged into PBS, and quantified by UV spectroscopy. The mammalian signal peptide MPLLLLLPLLWAGALA was used for both chains.

### 2.5 Quality control

Purified antibodies were analyzed by SDS-PAGE under reducing and non-reducing conditions. Purity and aggregation profile were assessed by SEC-HPLC and endotoxin level was measured by LAL chromogenic assay. Concentration, production yield, formulation buffer and physical inspection results were also reported.

### 2.6 Indirect ELISA binding assay

Microtitre plates were coated with the p95HER2-derived peptide antigen (MPIWKFPDEEGACQPCPINCTHSCVDLDDKGCPAEQRASPLTHHHHHH) at 2.5 µg/mL (50 µL/well) for 1 hour with shaking at 300 rpm and blocked overnight at 4 °C with 1% casein in PBS. The peptide was supplied as an unmodified TFA salt (lot U2097FD150-1/PE4504); supplier quality-control documentation reported 98.7% RP-HPLC purity and an ESI-MS mass of 5426.4 Da, consistent with the theoretical mass of 5427.05 Da. Antibodies were titrated in duplicate as a 3-fold dilution series starting at 3.0 µg/mL. Bound antibody was detected with HRP-anti-IgG and TMB substrate, stopped with 1 M HCl, and read at 450 nm.

### 2.7 Biacore kinetic analysis

Kinetics were measured on a Biacore T200. Antibodies were covalently immobilised by EDC/NHS chemistry and the peptide antigen was flowed as analyte in multi-cycle kinetics. The last three independent runs were globally fitted to a two-state reaction model to derive k_a1_, k_d1_, k_a2_, k_d2_ and the overall equilibrium dissociation constant K_D_.

## 3. Results

### 3.1 An integrated computational pipeline translates sequence data into interaction models to guide combinatorial variant generation

Starting from sequence alone, we modelled the 611-CTF-p95HER2 (p95HER2) ectodomain and the NAZ-mAb Fv and assembled the NAZ-mAb–p95HER2 complex by information-driven docking, taking the consensus of three docking methods (Dorraji et al. 2022) (**Figure 1A**). Using this complex as the wild-type reference, we defined the paratope as the antibody residues within 4.5 Å of p95HER2. These docking-derived interacting paratope residues agreed with our hydrogen-deuterium-exchange mass-spectrometry (HDX-MS) data (Dorraji et al. 2022). Therefore, using this complex as a wild type reference, we performed FoldX 5.0 (Delgado et al. 2019) saturation mutagenesis across the paratope, substituting each position with the remaining 19 amino acids and recording the change in interaction energy. The wild-type complex had interaction energy of −10.87 kcal/mol. 14 out of 23 positions in the paratope region showed destabilization in more than 10 point mutations, where interaction energy of the mutant was less compared to the wild-type (**Figure 1B**, **Table 1**). Among these, six positions (H:F38, H:W52, H:Y61, L:G33, L:K36, L:S114) had more than 17 out of 19 point mutations as destabilizing. These regions are potential binding hotspot regions, which might be essential for NAZ-mAb–p95HER2 binding.

**Figure 1.**
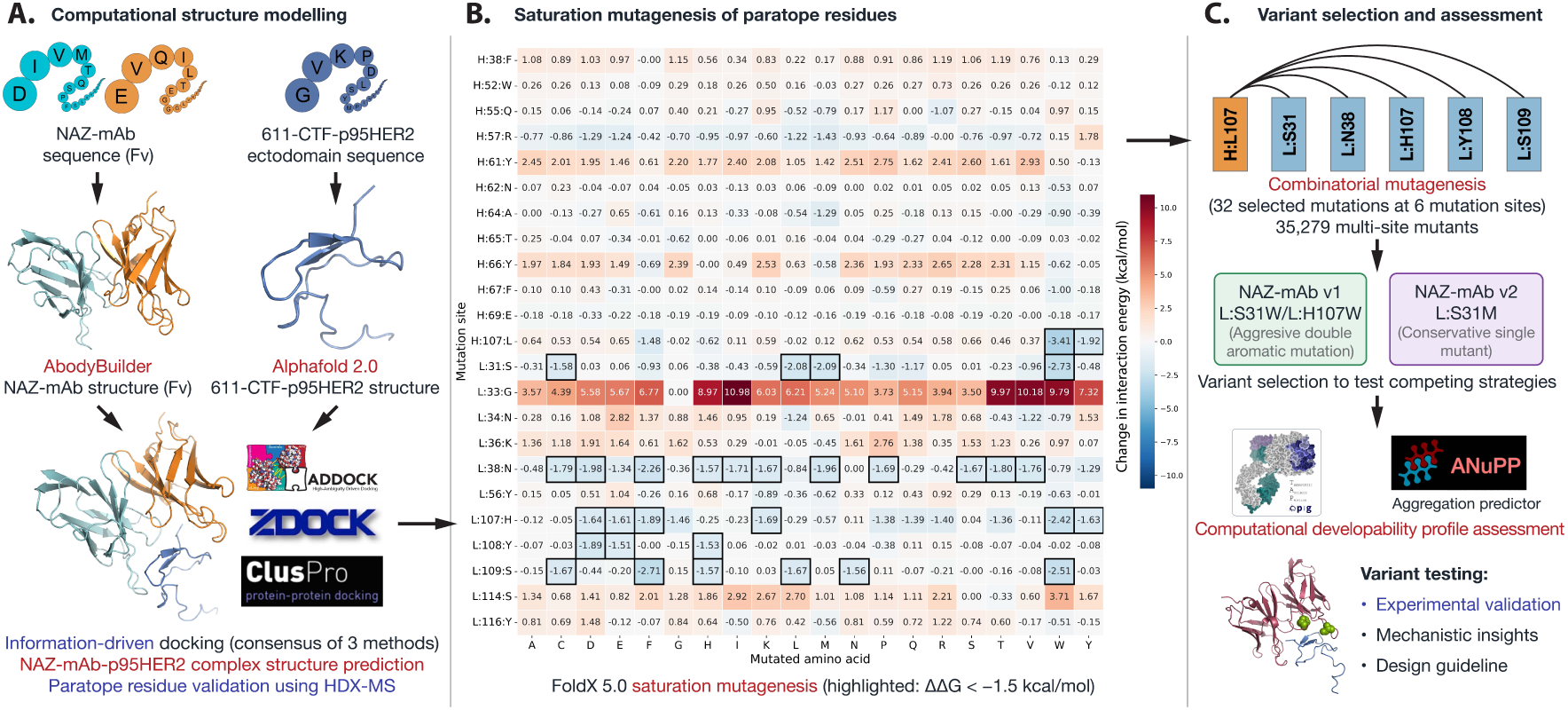
The schematic workflow of the computational study design. The starting points were the sequences of NAZ-mAb and 611-CTF-p95HER2 ectodomain. The computational pipeline includes modelling of the structure of both antibody and antigen, docking, mutational scanning of each paratope, combinatorial mutant generation and developability assessment. We selected two variants to test the computational variant design strategy and experimentally validated the results.

**Table 1.**
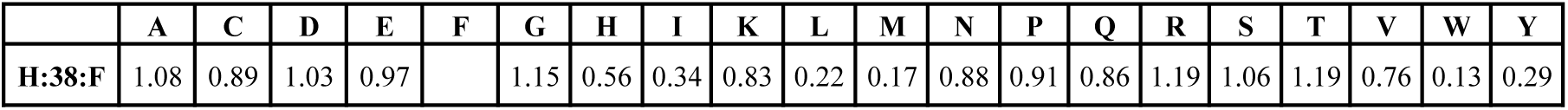
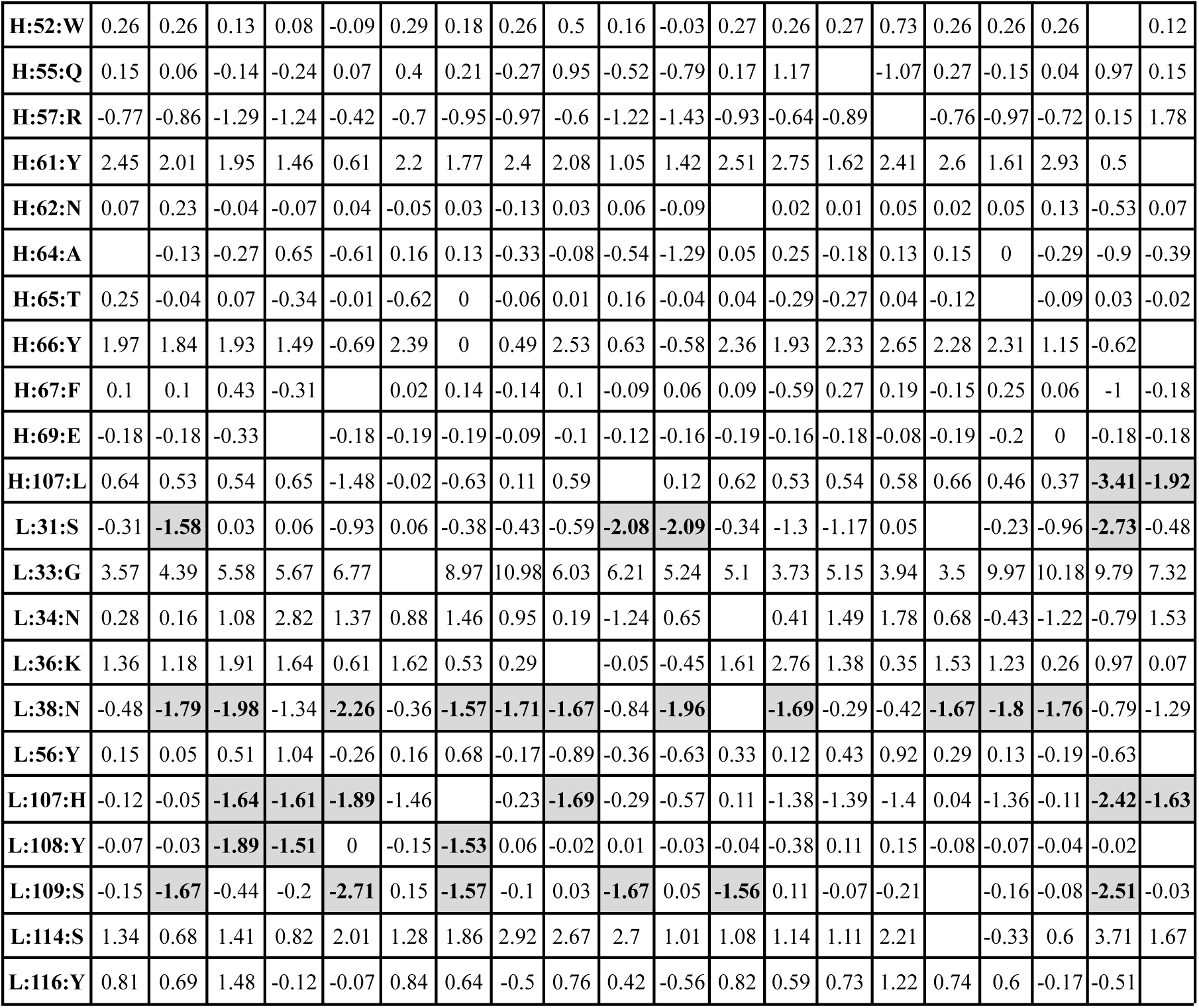
Change in interaction energy upon point mutations for all paratope residues in NAZ-mAb antibody. Highlighted mutations have change in interaction energy less than -1.5 kcal/mol at six mutation sites. Column headers denote the mutation and row headers denote mutation sites (chain:position:wild-type amino acid). Note that antibody residue positions are given in the IMGT numbering scheme. The wild-type interaction energy was -10.87 kcal/mol.

Moving from position to specific point mutation level, restricting attention to high-confidence binding-improving substitutions (Δ interaction energy < −1.5 kcal/mol) left six positions (heavy chain H:L107 and light-chain L:S31, L:N38, L:H107, L:Y108 and L:S109) and 32 qualifying point mutations. The most favourable point mutations were polar/non-polar-to-aromatic changes (**Figure 1B**, **Table 1**), the largest being heavy-chain H:L107W (Δ interaction energy= −3.41 kcal/mol). Interestingly, the only position identified in the CDRH3 region (H:L107) had two highly stabilizing aromatic substitutions (change in interaction energy ≤ -1.5 kcal/mol). However, the other 14 mutations destabilized the NAZ-mAb–p95HER2 complex.

We further focused on the 6 paratope sites containing 32 mutations (**Figure 1C**, **Table 2**) significantly improving binding affinity (≤ -1.5 kcal/mol), mainly to accommodate the error in computational calculation and reduce the computational time to prepare all mutation combinations (23^19^ possible combinations). For all combinatorial mutations (35,279 mutations; **Figure 2)**, the change in interaction energy was calculated using FoldX 5.0 (Delgado et al. 2019). We observed that change in interaction energies were normally distributed (**Figure 2B**), with almost all of the mutations improving the predicted interaction energy. Predicted interaction energy improved monotonically with the number of mutated sites from roughly −3 kcal/mol for the best single mutant to below −9 kcal/mol for the best six-site mutation (**Figure 2C**). It may reflect the largely additive manner in which FoldX 5.0 accumulates per-site contributions and does not account for the back-bone chain flexibility and stability due to the high mutational load. The favourable interaction energy for each mutational load (no. of site containing mutation) showed strong enrichment for tryptophan (H:L107W, L:S31W, L:H107W and L:S109W), where H:L107W was essentially obligatory in all high-ranking designs. These aromatic mutations were also high interaction energy mutants in our previous saturation mutagenesis experiment as well.

**Figure 2.**
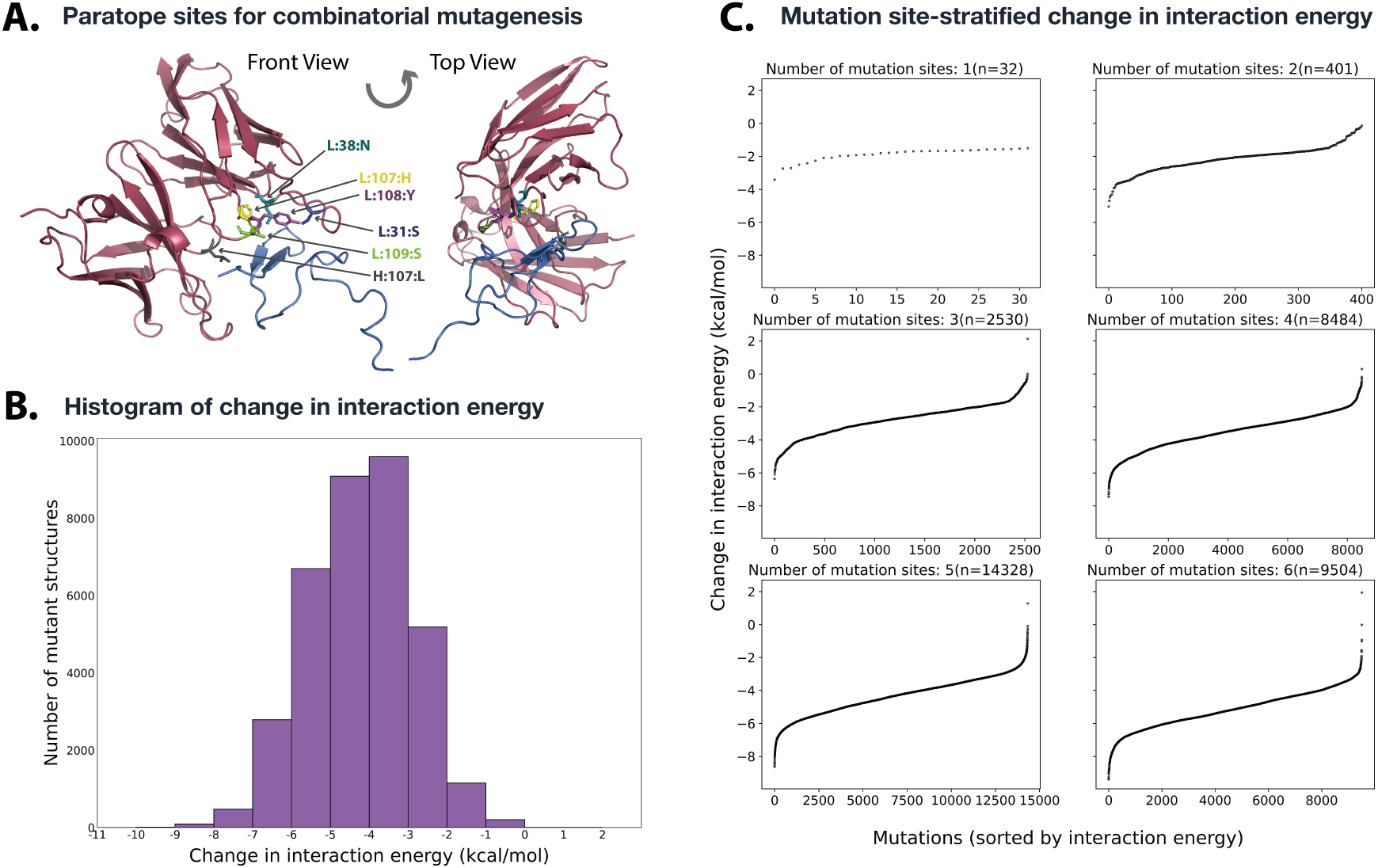
Combinatorial mutagenesis of NAZ-mAb provides variant panels to test the design strategy. **(A)** The six mutation sites, selected by saturation mutagenesis (mutating with remaining 19 amino acids) of paratope residues, are shown in the structure of the NAZ-mAb–611-CTF-p95HER2 complex as a front view (left) and top view (right). **(B)** the distribution of the change in interaction energy shown as a histogram for all combinatorial mutations, where most of the mutations show additive nature for multiple mutants and a very small percent of mutants have worse interaction energy than the wild-type. **(C)** The sorted distribution of change in interaction energy segregated by no. of mutation sites.

**Table 2.**
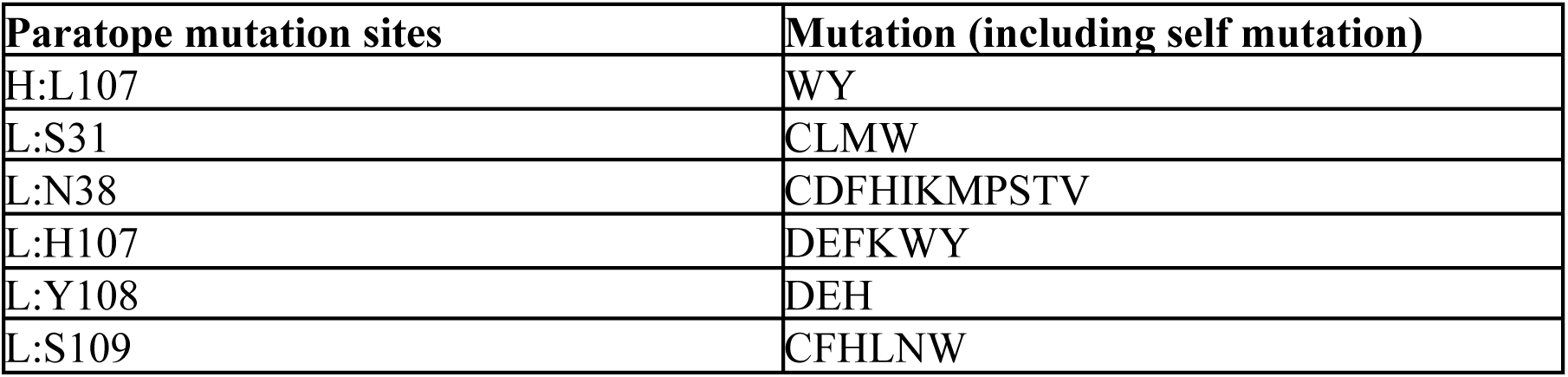
The selected mutations for the combinatorial mutagenesis in up to six positions. A total of 35,279 multi-order mutations were generated. Paratope sites are denoted as chain:position:wild-type amino acid.

### 3.2 Computational selection of NAZ-mAb variants to evaluate conservative versus aggressive rational design strategies

To provide a mechanistic understanding to design better variants, we carefully selected two variants for the experimental validation as follows: (i) selected an aromatic hotspot combination that tests whether combining recurrent high-impact sequentially distant aromatic substitutions improves interaction at the cost of larger physicochemical perturbation (NAZ-mAb v1; L:S31W/L:H107W), (ii) test whether a single conservative, non-aromatic light chain mutation can preserve the scaffold while modestly improving binding (NAZ-mAb v2; L:S31M). We avoided the site H:L107 in the CDRH3 region, since most mutations at this site were destabilizing except for the aromatic mutation Trp (W) and Tyr (Y) (**Table 1**). An additional variant bearing five light chain substitutions (L:S31W/L:N38M/L:H107W/L:Y108D/L:S109W) was also designed with significantly improved interaction energy; however, no detectable expression was obtained, likely reflecting the cumulative destabilizing effect of the mutations.

First, we computationally assessed the developability profile of the NAZ-mAb antibody variant with respect to wild-type and other clinically approved therapeutic antibodies using Therapeutic antibody profiler (TAP, https://opig.stats.ox.ac.uk/webapps/sabdab-sabpred/sabpred/tap) (Raybould et al. 2024). It compares the developability profile of the antibody of interest with that of approved therapeutic antibodies by comparing five biophysical features: (i) Total CDR length, (ii) Patches of surface hydrophobicity (PSH), (iii) Patches of Positive Charge (PPC), (iv) Patches of Negative Charge (PNC) and (v) Structural Fv Charge Symmetry Parameter (SFvCSP) (**Table S1)**. All parameters fell not only within the developability envelope but within the highest-density region of the latest set of 754 post-Phase-1 therapeutic Fv domains (**Figure S1, Table 3**). As expected, the developability profile of NAZ-mAb v2 (L:S31M) was closest to the wild-type compared to the NAZ-mAb v1 (L:S31W/L:H107W). Additionally, we predicted aggregation-prone regions in the NAZ-mAb antibody and its variants using ANuPP (Prabakaran et al. 2021). At the default threshold 0.52, no APR was observed at this region, highlighting low aggregation propensity of the wild-type and its variants.

**Table 3.** Developability profile of NAZ-mAb antibody variants calculated from Therapeutic antibody profiler (TAP) webserver. Amber/red guideline thresholds are given in **Table S1**.

| Metric | Wild-type | L:S31W/L:H107W | L:S31M |
| --- | --- | --- | --- |
| Total CDR Length | 49 | 49 | 49 |
| CDR Vicinity PSH Score (Kyte & Doolittle) | 143.123 | 145.4371 | 142.1395 |
| CDR Vicinity PPC Score | 0.1451 | 0.1067 | 0.158 |
| CDR Vicinity PNC Score | 0.0515 | 0.1386 | 0.0531 |
| SFvCSP Score | 0 | -1 | -0.9 |

### 3.3 NAZ-mAb v2 shows a marked recombinant expression-yield advantage over parental NAZ-mAb and v1

Parental NAZ-mAb and both engineered variants were produced as mouse-IgG1 antibodies at 1 mg/mL in PBS. The variants were reported as >98% monomer by SEC-HPLC, had low endotoxin levels, and showed the expected SDS-PAGE patterns under reducing and non-reducing conditions (**Table 4**). Expression yield differed substantially among the three antibodies. NAZ-mAb v2 yielded 37.5 mg/L, compared with 7.5 mg/L for parental NAZ-mAb and 6.25 mg/L for NAZ-mAb v1. Thus, the single L:S31M substitution increased recombinant yield five-fold relative to parental NAZ-mAb and six-fold relative to v1, whereas v1 was produced at a yield similar to parental NAZ-mAb.

**Table 4.**
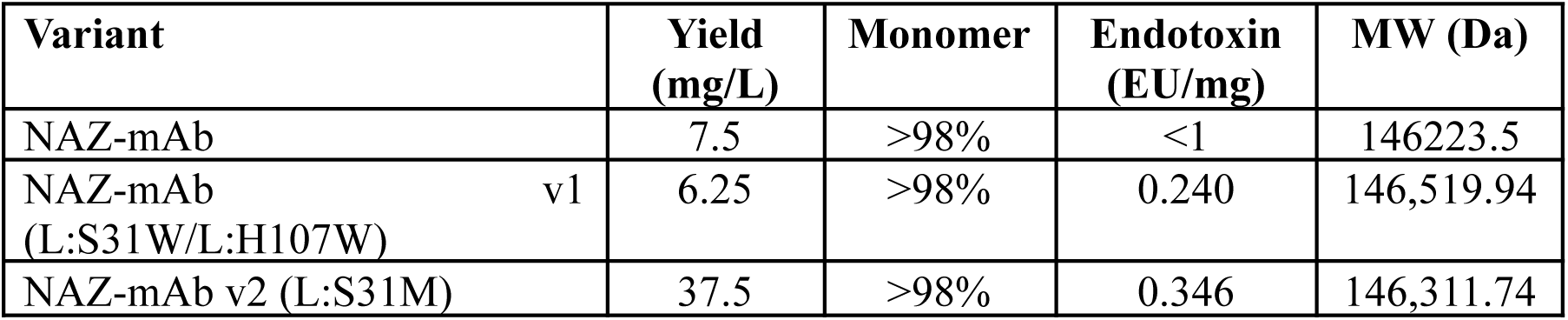
Production and yield summary of the NAZ-mAb antibody variant.

| Variant | Yield (mg/L) | Monomer | Endotoxin (EU/mg) | MW (Da) |
| --- | --- | --- | --- | --- |
| NAZ-mAb | 7.5 | >98% | <1 | 146223.5 |
| NAZ-mAb v1 (L:S31W/L:H107W) | 6.25 | >98% | 0.240 | 146,519.94 |
| NAZ-mAb v2 (L:S31M) | 37.5 | >98% | 0.346 | 146,311.74 |

### 3.4 Engineered NAZ-mAb variants exhibit enhanced apparent binding in ELISA but weaker monovalent affinities in Biacore analysis

In indirect ELISA both engineered variants produced higher absorbance than parental NAZ-mAb across the titration, with NAZ-mAb v2 highest and NAZ-mAb v1 matching or exceeding parental at higher concentrations (**Fig.3B**). Taken alone, this would suggest improved binding. However, Biacore two-state kinetics inverted the ranking (**Table 5**), where parental NAZ-mAb had the best K_D_ at 5.33 nM, NAZ-mAb v2 was close behind at 9.45 nM, and NAZ-mAb v1 was weakest at 32.6 nM (6.1-fold loss relative to parental). The NAZ-mAb v1 deficit was driven by both a slower on-rate and a faster off-rates (**Table 5**). Critically, the experimental affinity order is the reverse of the FoldX 5.0 prediction, in which NAZ-mAb v1 (-14.2684 kcal/mol) was predicted to bind more tightly than NAZ-mAb v2 (-12.96 kcal/mol).

**Figure 3.**
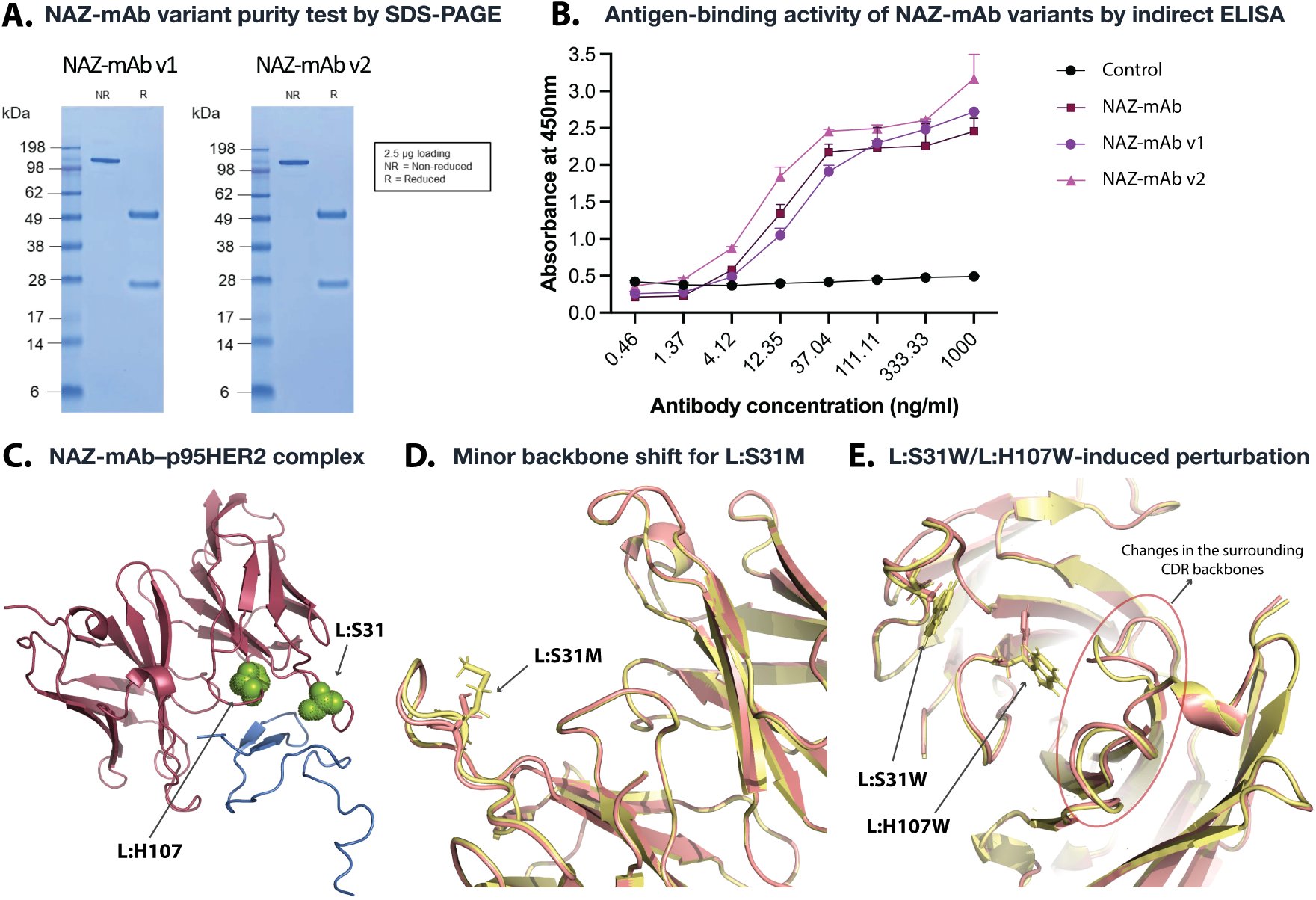
Experimental validation and structural assessment of the variants. **(A)** SDS-PAGE analysis of purified NAZ-mAb v1 and NAZ-mAb v2 under non-reduced (single band, >100 kDa) and reduced (two bands, ∼50 kDa and ∼25–28 kDa) conditions (2.5 µg loading), confirm correct assembly and purity of the expressed variants. **(B)** ELISA dose-response curves showing antigen-binding activity of NAZ-mAb (wild-type), NAZ-mAb v1, and NAZ-mAb v2 against 611-CTF-p95HER2 across a concentration range of 0.46–1000 ng/ml. Absorbance was measured at 450 nm. Both variants produced higher apparent peptide-binding signals than parental NAZ-mAb in the indirect ELISA format, with NAZ-mAb v2 generating the highest signal at the upper antibody concentrations. The control shows no binding. The FoldX 5.0 mutated structure does not account for the variation in the backbone chain upon mutations. **(C)** The structure of the wild-type NAZ-mAb (red) in complex with 611-CTF-p95HER2 ectodomain (blue), where mutation sites in the light chain are highlighted as green spheres. **(D)** The change in interaction interface upon conservative mutations L:S31M (highlighted as stick representation) leads to minor change only in CDRL1 backbone. **(E)** The change in interaction interface upon double aromatic mutations (L:S31W/L:H107W) also shows minor variation at the CDRL1 backbone. However, these double mutations alter the structure of CDRH2 and CDRH3 significantly (especially at position H:L107 and H:R108). The wild-type structure is in light red color and mutated structure is in light yellow color for **(D)** and **(E)**.

**Table 5.** Biacore two-state kinetics (mean of three runs). The fold change is given with respect to the wild-type.

| Antibody | $k_{a1}$ ($M^{-1}s^{-1}$) | $k_{d1}$ ($s^{-1}$) | $k_{a2}$ ($s^{-1}$) | $k_{d2}$ ($s^{-1}$) | $K_D$ (nM) | Fold change |
| --- | --- | --- | --- | --- | --- | --- |
| NAZ-mAb | $5.07 \times 10^4$ | $8.06 \times 10^{-3}$ | $3.46 \times 10^{-3}$ | $1.75 \times 10^{-3}$ | 5.33 | |
| NAZ-mAb v1 | $1.61 \times 10^4$ | $3.29 \times 10^{-2}$ | $1.62 \times 10^{-2}$ | $3.08 \times 10^{-3}$ | 32.6 | $6.1 \times$ |
| NAZ-mAb v2 | $3.40 \times 10^4$ | $5.75 \times 10^{-2}$ | $1.23 \times 10^{-2}$ | $7.25 \times 10^{-4}$ | 9.45 | $1.8 \times$ |

### 3.5 State-of-the-art computational tools show inconsistent predictions for variant stability and affinity

We also used other tools to assess the binding affinity and stability of the variants with respect to the wild-type. The MutaBind2 (Zhang et al. 2020) predicted minor change in binding affinity induced by a mutation for NAZ-mAb v1 (-0.06 kcal/mol) and de-stabilization of binding for NAZ-mAb v2 (0.54 kcal/mol). The stabilization for the aromatic double mutant is aligned with the FoldX 5.0 prediction, highlighting potential biases for aromatic residues. To further assess the stability of the complex upon mutation, we estimated the change in stability upon mutations for the complex with FoldX 5.0 (Delgado et al. 2019) and DynaMut2 (Rodrigues et al. 2021). FoldX 5.0 predicted both mutations as more stable than the wild-type structure (L:S31M ΔΔG = -1.25 kcal/mol and L:S31W/L:H107W ΔΔG = -1.32 kcal/mol), aligned with the predicted affinity. This observation aligns with a previous study that showed that most affinity prediction tools are basically learning the stability prediction (de Kanter et al. 2026). On the other hand, DynaMut2 predicted stabilization for v2 L:S31M (ΔΔG = 0.18 kcal/mol) and destabilization for v1 L:S31W/L:H107W (ΔΔG = -0.39 kcal/mol). DynaMut2 uses opposite signs (positive ΔΔG is stabilizing) and only predicts the stability of the complex. The observation from different methods shows that there was no alignment in the change in binding affinity across methods. However, all tools predicted improvement in stability for the NAZ-mAb v2.

### 3.6 Bulky aromatic mutations induce significant CDR backbone rearrangements that potentially compromise binding affinity

To validate that FoldX 5.0 showed predicted affinity was off due to limitation of FoldX 5.0 not correcting for the changes in the backbone, we modelled the structure of the NAZ-mAb variants using Abodybuilder (**Fig.3C**). The backbone of modelled structure of the conservative mutation (NAZ-mAb v2) was almost perfectly aligned with the wild-type with minor changes at the site of mutation (**Fig. 3D**). For the double aromatic mutation (NAZ-mAb v1), the variation at the mutation site was also negligible. However, due to the large aromatic mutations at the interface, it changed the conformation of the CDRH2 and CDRH3, which is likely reflected in the higher change in K_D_ observed in the kinetic experiments (**Fig. 3E**). Two residues at the CDRH3 H:L107 and H:R108 had significant rearrangement in their 3D position.

## 4. Discussion

The entire pipeline began from sequence alone; the 611-CTF-p95HER2 ectodomain and NAZ-mAb Fv structures were modelled computationally. In the absence of an experimental co-structure, the antibody–antigen complex was assembled by information-driven docking, using the experimentally-validated “MPIWKFPDEE” region in the p95HER2 extracellular domain as the potential epitope region. This single, computationally generated complex served as the wild-type reference for all subsequent energetic calculations. The antibody–antigen interaction was dominated by CDRH2 and CDRL3 residues, consistent with prior functional evidence from HDX-MS experiments (Dorraji et al. 2022). FoldX 5.0 saturation scanning identified core interaction hotspots in CDRH2 that are essential for retaining binding. By contrast, substitutions in CDRL3 showed improvement in predicted binding affinity and were therefore selected as sites to engineer novel NAZ-mAb variants.

FoldX 5.0 is an empirical force field parameterised principally to predict the stability effect (ΔΔG of folding) of point mutations, computed on an essentially fixed backbone using static van der Waals packing, hydrogen bonding, and a mean-field solvation term (Delgado et al. 2019). To design novel variants and gain mechanistic insight, we first pursued an aggressive strategy by introducing two aromatic substitutions (NAZ-mAb v1, L:S31W/L:H107W), replacing a small polar residue (Ser) and a charged residue (His) with large aromatic side chains. A rigid backbone rewards new aromatic substitutions through favourable packing and π-contacts but cannot account for the corresponding costs; the conformational entropy of immobilising a bulky indole ring in the bound state, the desolvation penalty, and the backbone strain required to accommodate the larger side chains. Benchmarking of ΔΔG methods has demonstrated this failure mode, where ensemble/backbone-sampling approaches outperform fixed-backbone scoring most strongly for small-to-large side-chain mutations and for multiple simultaneous non-alanine mutations (Sanavia et al. 2020; Potapov et al. 2009; Barlow et al. 2018). Furthermore, the two aromatic mutations lie in close proximity and may sterically compete for the same interface pocket, co-distorting the loop geometry in ways that a rigid-backbone model cannot capture. As a result, the effect of multiple mutations that are likely to change the whole or part of the interface is not truly captured. Moreover, aromatic residues and proline are overestimated in the FoldX 5.0 observations (Vliora et al. 2026). This limitation is directly reflected in our data: although FoldX 5.0 predicted a substantial improvement in interaction energy for the double aromatic mutant (−14.27 kcal/mol versus −10.87 kcal/mol for wild-type), the experimentally measured K_D_ was 6.1-fold weaker than parental (32.6 nM versus 5.33 nM).

Our more conservative design NAZ-mAb v2 (L:S31M), stays inside the regime where fixed-backbone prediction is dependable and where we avoided mutation with aromatic residue. The residues in the light-chain position were mutated where most substitutions seemed to improve performance, leading to the hypothesis that this position is not essential for binding. The mutated single residue was non-polar and hydrophobic (M). Although FoldX 5.0 showed a slight improvement in predicted binding affinity (dG = -12.9574 kcal/mol), the K_D_ showed a slight decrease (9.45 nM). In comparison to these mutants, the interaction energy of the wild-type was -10.87 kcal/mol and the K_D_ was 5.33 nM. This conservative single mutant nearly retained parental affinity compared to the double aromatic mutants (1.8-fold versus 6.1-fold loss). The data clearly illustrate that, for this paratope, FoldX 5.0 interaction-energy ranking is a usable hotspot-identification filter but not a quantitative affinity predictor for combinatorial designs. Although FoldX 5.0 was not able to identify the mutants that improved the K_D_, we obtained mechanistic insights into how to identify such mutants given the limitations of empirical energy-based tools.

Although the two engineered variants share the same heavy chain, their recombinant expression yields differed substantially. NAZ-mAb v2 achieved a yield of 37.5 mg/L, representing a five-fold increase over parental NAZ-mAb (7.5 mg/L) and a six-fold increase over NAZ-mAb v1 (6.25 mg/L). In contrast, v1 was produced at a yield broadly comparable to the parental antibody. These results identify v2 as the strongest production variant under the tested expression conditions, consistent with predictions from two independent stability prediction tools (FoldX 5.0 and DynaMut2). The surface-hydrophobicity score from the TAP profile decreased only marginally for v2 relative to parental NAZ-mAb (PSH 143.1 to 142.1), with both values remaining below the TAP amber threshold of 151.1. While these observations do not establish a relationship between stability or PSH and recombinant yield due to the limited dataset size, they suggest the potential for further investigation using a larger mutant dataset. ANuPP predicted no aggregation-prone regions in parental NAZ-mAb or either engineered variant. Consistent with this, both purified variants were reported as >98% monomer by SEC-HPLC and had low endotoxin levels.

Interestingly, ELISA ranks the variants above parental, Biacore ranks them below. The different rank orders are consistent with the fact that the two assays measure binding under distinct antigen-presentation and assay geometries. Indirect ELISA uses a plate-immobilised peptide and a bivalent IgG, so its signal convolves intrinsic affinity with avidity (bivalent re-binding to densely coated antigen), coating density and detection gain, and it plateaus where avidity and antigen capacity saturate. However, the peptide also contains five cysteines and, although supplier quality-control data confirmed high purity and an intact mass consistent with the expected sequence, its cysteine oxidation state, free-thiol content and disulfide connectivity were not directly characterised. Whereas Biacore measures the monovalent K_D_. Avidity routinely masks several-fold differences in monovalent affinity, so a higher ELISA absorbance for a bivalent reagent is not evidence of tighter intrinsic binding. Biacore, with immobilised antibody and peptide analyte fitted to a two-state model, isolates the monovalent kinetic *K_D_* and is the appropriate arbiter of affinity. Therefore, the variants show enhanced apparent peptide binding in the ELISA format, particularly NAZ-mAb v2, but the higher ELISA signal does not by itself establish improved intrinsic affinity.

## 5. Limitations

The most fundamental limitation is that the entire energetic analysis rests on a computationally generated, rather than experimentally determined, structure. Absolute interaction energies must therefore be treated as relative guides only, and the affinity inversion is in part attributable to compounding model error at an unverified interface. The docking-derived epitope agreed with HDX data and the hotspot map with prior functional evidence, which lends confidence to the interface, but a co-crystal or cryo-EM structure would be required to ground the energetics quantitatively. Standard MD were not able to provide decisive ranking power in this system because the antigen was highly flexible, making the conformational space too large to sample adequately within realistic simulation time and across replicate. Beyond the structural basis, the two produced variants share a heavy chain and sample only two points of a large design space; The Biacore *K_D_* values derive from a two-state fit whose micro-rate constants are model-dependent, although the overall *K_D_* ranking is robust. Finally, all binding reported here is to a short synthetic peptide. Although supplier quality-control data confirmed high peptide purity and an intact mass consistent with the expected sequence, cysteine oxidation state and disulfide connectivity were not directly characterised and may have influenced peptide presentation across assay formats. Whether the engineered variants retain engagement of full-length, membrane-embedded 611-CTF-p95HER2 in a cellular context remains to be established.

## 6. Conclusion

This study evaluated a sequence-to-structure computational workflow for engineering NAZ-mAb variants against the 611-CTF-p95HER2 target. The workflow generated experimentally tractable candidates and distinguished between two contrasting design strategies: a conservative single substitution, NAZ-mAb v2 (L:S31M), and an aromatic double substitution, NAZ-mAb v1 (L:S31W/L:H107W). Although neither variant improved the measured Biacore affinity relative to parental NAZ-mAb, both retained p95HER2 engagement and showed enhanced apparent peptide binding in indirect ELISA, with NAZ-mAb v2 producing the highest ELISA signal. This improved performance in the plate-based assay does not, by itself, demonstrate improved intrinsic affinity relative to the parental antibody.

NAZ-mAb v2 emerged as the best-balanced engineered variant. It retained low-nanomolar p95HER2 binding by Biacore, showed enhanced apparent peptide binding in ELISA, and achieved a recombinant expression yield of 37.5 mg/L—five-fold higher than parental NAZ-mAb (7.5 mg/L) and six-fold higher than NAZ-mAb v1 (6.25 mg/L). This substantial production advantage was achieved via improving the stability of the complex, while retaining a TAP developability profile closely comparable to parental NAZ-mAb. In addition, no aggregation-prone regions were predicted for parental NAZ-mAb or either variant, and both purified variants were reported as >98% monomer by SEC-HPLC under the tested conditions.

Collectively, these findings show that fixed-backbone interaction-energy ranking alone is insufficient to predict experimentally measured kinetic affinity for higher-order mutants. For the NAZ-mAb scaffold, the conservative L:S31M substitution provided a more productive engineering solution than stacking additional aromatic substitutions. It preserved strong target engagement, enhanced apparent binding in the ELISA format, and delivered a marked manufacturability advantage. NAZ-mAb v2 therefore represents the strongest lead from this design set for further development of p95HER2-targeting molecules.

