## Supplementary Material for "Structure-guided computational design and mechanistic understanding of the p95HER2-targeting NAZ-mAb antibody and its variants"

* Corresponding author

Email:

**Supplementary Figure S1.** The developability profile of NAZ-mAb and its variants (green color) with respect to the 754 post-Phase-1 therapeutic Fv domains (shown as a histogram). The orange and red colors show the amber flag and red flag region, respectively, as defined by the Therapeutic antibody profiler (TAP).

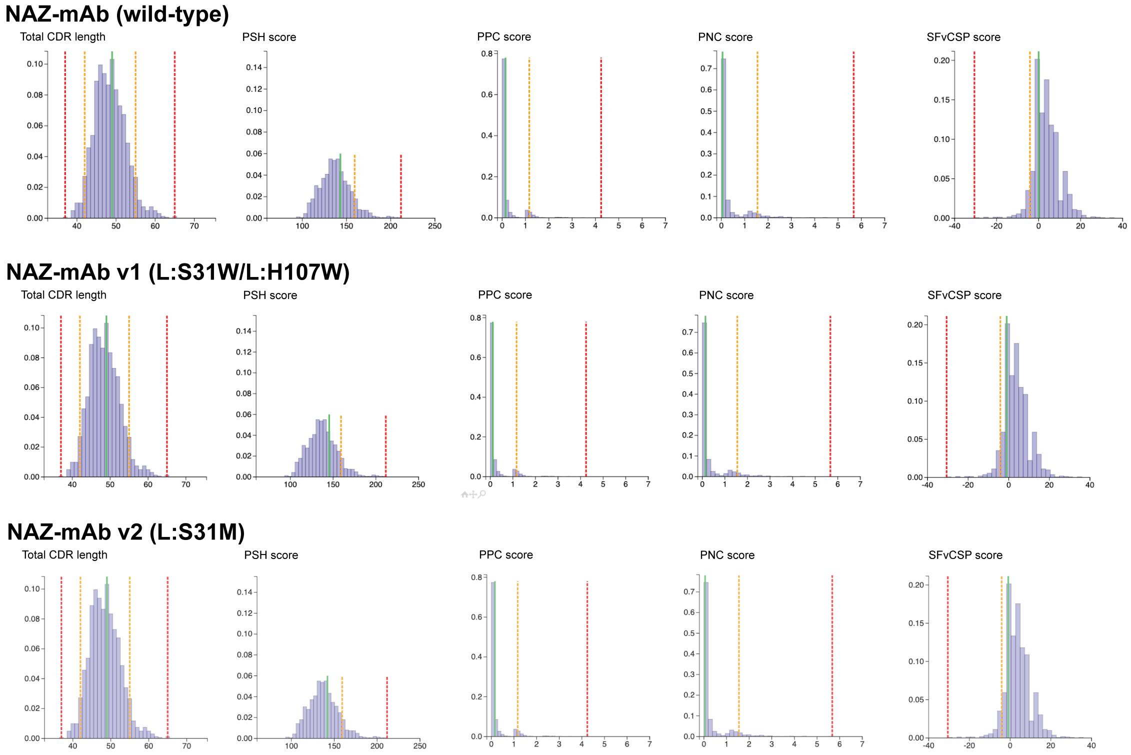

**Table S1.** The cutoffs for the developability parameters taken from the Therapeutic antibody profiler (TAP) webserver.

| **Developability parameters** | **Red flag region cutoff** | **Amber flag region cutoff** |
|---|---|---|
| **Total CDR length** | <38 and 63< | <43 and 55< |
| **Patches of surface hydrophobicity (PSH)** | <77.6 to 182.5< | <94.9 to 151.1< |
| **Patches of Positive Charge (PPC)** | >3.71 | >1.3 |
| **Patches of Negative Charge (PNC)** | >3.65 | >1.97 |
| **Structural Fv Charge Symmetry Parameter (SFvCSP)** | <-19.59 | <-4.95 |
